# Going down the rabbit hole: a review on methods characterizing selection and demography in natural populations

**DOI:** 10.1101/052761

**Authors:** Yann X.C. Bourgeois, Khaled M. Hazzouri, Ben H. Warren

## Abstract

1. Characterizing species history and identifying loci underlying local adaptation is crucial in functional ecology, evolutionary biology, conservation and agronomy. The ongoing and constant improvement of next-generation sequencing (NGS) techniques has facilitated the production of an ever-increasing number of genetic markers across genomes of non-model species.
2. The study of variation in these markers across natural populations has deepened the understanding of how population history and selection act on genomes. Population genomics now provides tools to better integrate selection into a historical framework, and take into account selection when reconstructing demographic history. However, this improvement has come with a burst of analytical tools that can confuse users.
3. Such confusion can limit the amount of information effectively retrieved from complex genomic datasets. In addition, the lack of a unified analytical pipeline impairs the diffusion of the most recent analytical tools into fields like conservation biology.
4. To address this need, we describe possible analytical protocols and link these with more than 70 methods dealing with genome-scale datasets. We summarise the strategies they use to infer demographic history and selection, and discuss some of their limitations. A website listing these methods is available at www.methodspopgen.com.

## Introduction

Multiple historical and selective factors shape the genetic makeup of populations. The advent of Next-Generation Sequencing (NGS) in the last 10 years has enhanced our understanding on how intermingled these factors are, and how they can impact genomic variation. Important results have been gathered on model species, or species of economic interest. Such results include, among other examples, an improved understanding of the history of human migrations, admixture and adaptation (e.g. Sabeti *et al.*, 2002; Abi-Rached *et al.*, 2011; Li and Durbin, 2011), the origin of domesticated species (e.g. Axelsson *et al.*, 2013; Schubert *et al.*, 2014), and the genetic basis of local adaptation in both model and non-model species (e.g. Legrand *et al.*, 2009; Kolaczkowski *et al.*, 2011; Roux *et al.*, 2013; Kubota *et al.*, 2015). The amount of population genomic data that is aimed at elucidating the history of natural populations has increased enormously in the last five years, even for non-model species. Studying genetic variation at the genome level allows the demographic factors shaping species history to be characterised. Further, understanding demographic history is important in correctly identifying loci under selection. Such data can even help in conservation efforts by identifying locally adapted genes that can be used to define relevant conservation units (Fraser and Bernatchez, 2001).

In the last 10 years, developments in NGS have continually improved the throughput of data, while reducing time and cost of their production. These methods have become more affordable for teams studying evolutionary processes in biology, and many new methods to infer demography and selection have been developed. However, these methodological advances have brought increased analytical complexity to the field, and an inflation in the number of methods covering any one topic. As a consequence, it has become increasingly difficult for all potential users to follow developments and be sure of selecting the most appropriate method for the question and data in hand.

An overarching theme that concerns new users in a wide range of contexts is understanding patterns of heterogenous diversity along the genome. Patterns of nucleotide variation in genomes are shaped by both intrinsinc and extrinsic factors. Even within a single isolated panmictic population, interaction between recombination, selection and historical variation in population size will lead to heterogeneous diversity along the genome. At the scale of several connected populations or even between emerging species, these processes will affect the rate at which migration homogenizes the genome (Wolf and Ellegren, 2016).

A prime example is the situation of a researcher primarily interested in identifying signatures of recent positive selection in a species of interest. Since a new mutation will see its frequency increase in a population where it provides a selective advantage (*i.e.* hard selective sweep), a large region around it can remain uniform, especially if selection is strong (Sabeti *et al.*, 2002; McVean, 2007; Vitti *et al.*, 2013). This can lead to an increase in linkage disequilibrium (LD) between variants associated to the advantageous mutation, as well as a decrease in the age of the positively selected alleles and their nucleotide diversity. If positive selection occurs only in some populations, it may be possible to observe an increase in differentiation at this locus (Charlesworth *et al.*, 1997). To detect this signature of selection, some methods can track particularly long haplotypes and linkage disequilibrium along the genome. Others will rather focus on allele frequency spectrum and nucleotide diversity. Association methods will take advantage of preliminary knowledge of a phenotype or environment to identify loci displaying correlated allele frequencies. A few methods aim at inferring the whole history of coalescence and recombination along genomes, but still make simplifying assumptions and often require whole-genome resequencing data, which remain unaffordable for many teams.

Therefore, the choice of methods of any such researcher will depend on the available data and specifics of the question being addressed. One key aspect is that all these methods and questions do not have the same requirements in terms of reference genomes and marker density. For example, recent discussion of RAD-markers has been interesting from this perspective (Lowry *et al.*, 2016; Catchen *et al.*, 2017). The density of markers obtained along a genome depends on the choice of the restriction enzyme, and this choice must take into account the average extent of LD. Genome scans of selection will lose power if this density is not enough to cover mutations in strong linkage with variants under selection.

In the absence of any unified framework, combining several tools is necessary to interpret results. It must be borne in mind that recombination rates vary along the genome, which can possibly bias tests based on LD. It can therefore be important to characterize the recombination landscape in natural populations, requiring the use of another method (e.g. LDHat, Table 1). Background selection can lead to signatures of high differentiation that mimick disruptive selection (Charlesworth *et al.*, 1997). An assessment of genetic diversity within populations, haplotype frequencies and possibly association with phenotype in each population would therefore be needed to explore this possibility (Charlesworth *et al.*, 1997). Demographic history impacts patterns of LD, allele age and frequencies at the genome scale, and affects the efficiency of selection at specific genes. This calls for at least basic checking of demographic structure and history and ideally building neutral demographic models to estimate the expected frequency of outliers without involving selection. In addition, most methods estimating selection coefficients require estimating effective population sizes. Finally, including markers under selection can bias demographic inference by skewing allele frequency spectra and LD, which requires careful data filtering and removal of outliers.

**Table 1.**
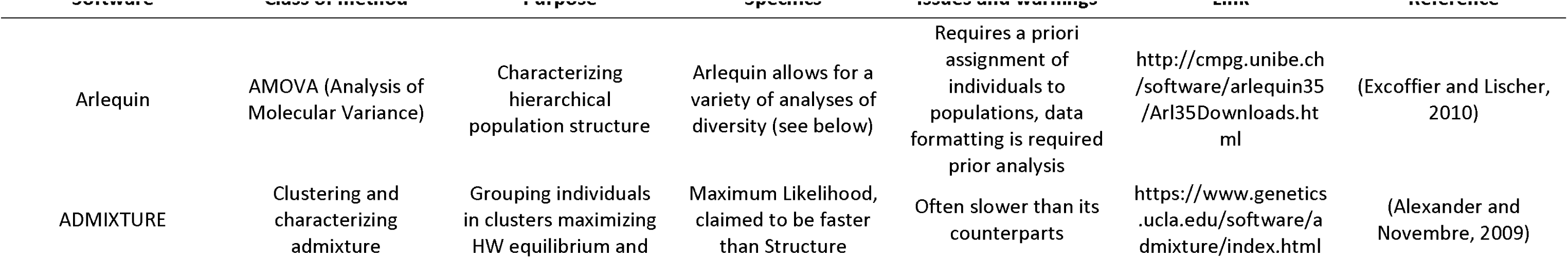
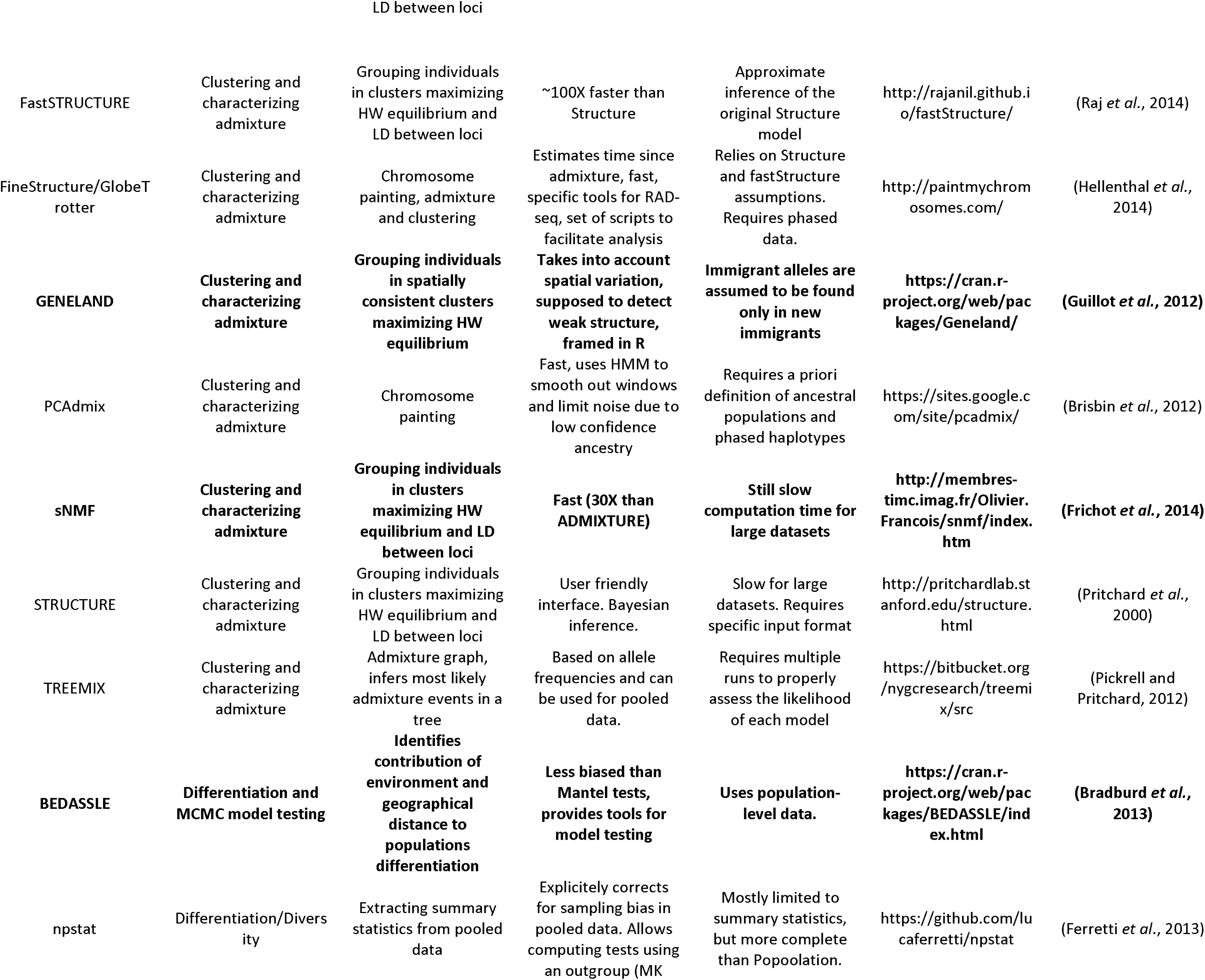
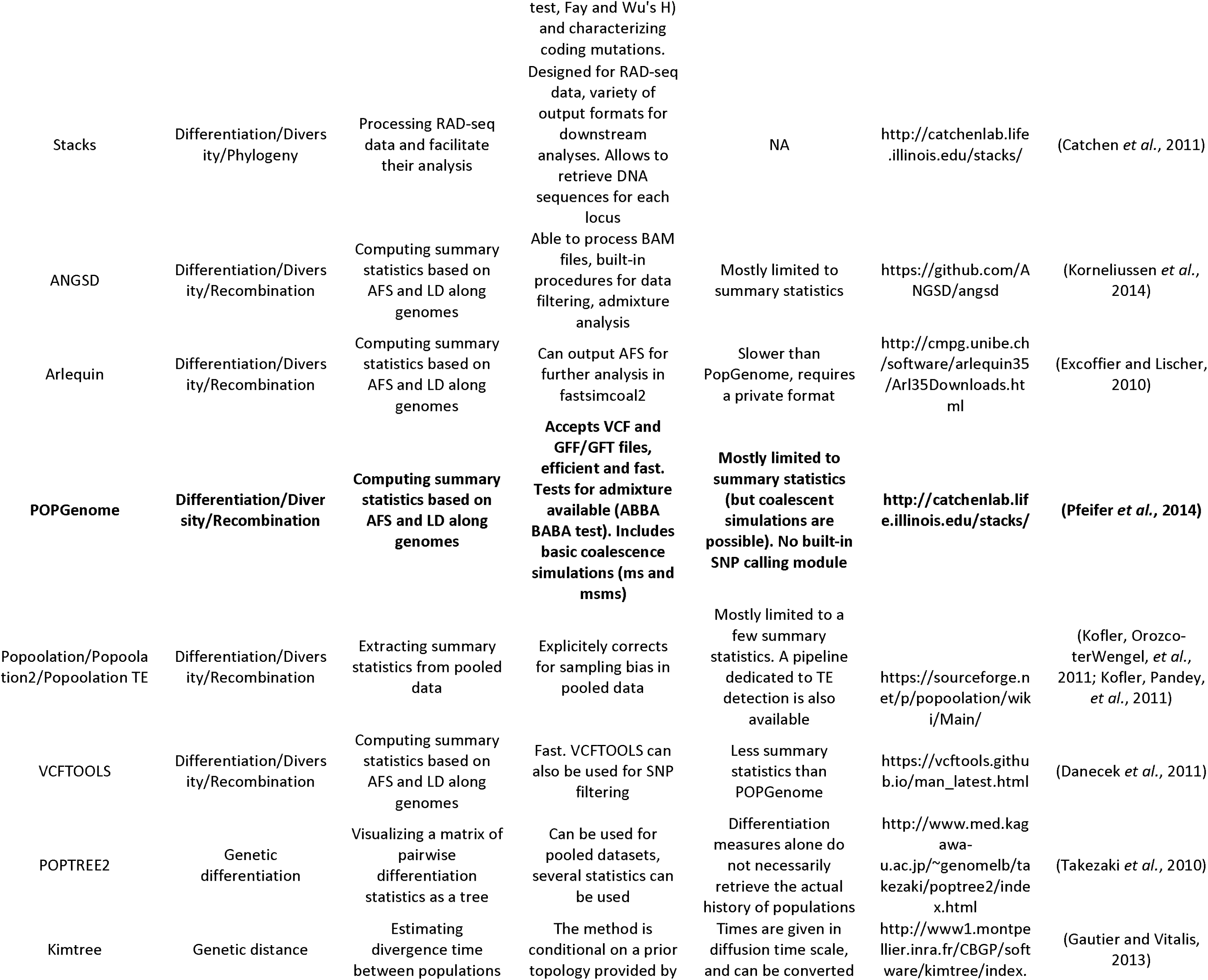
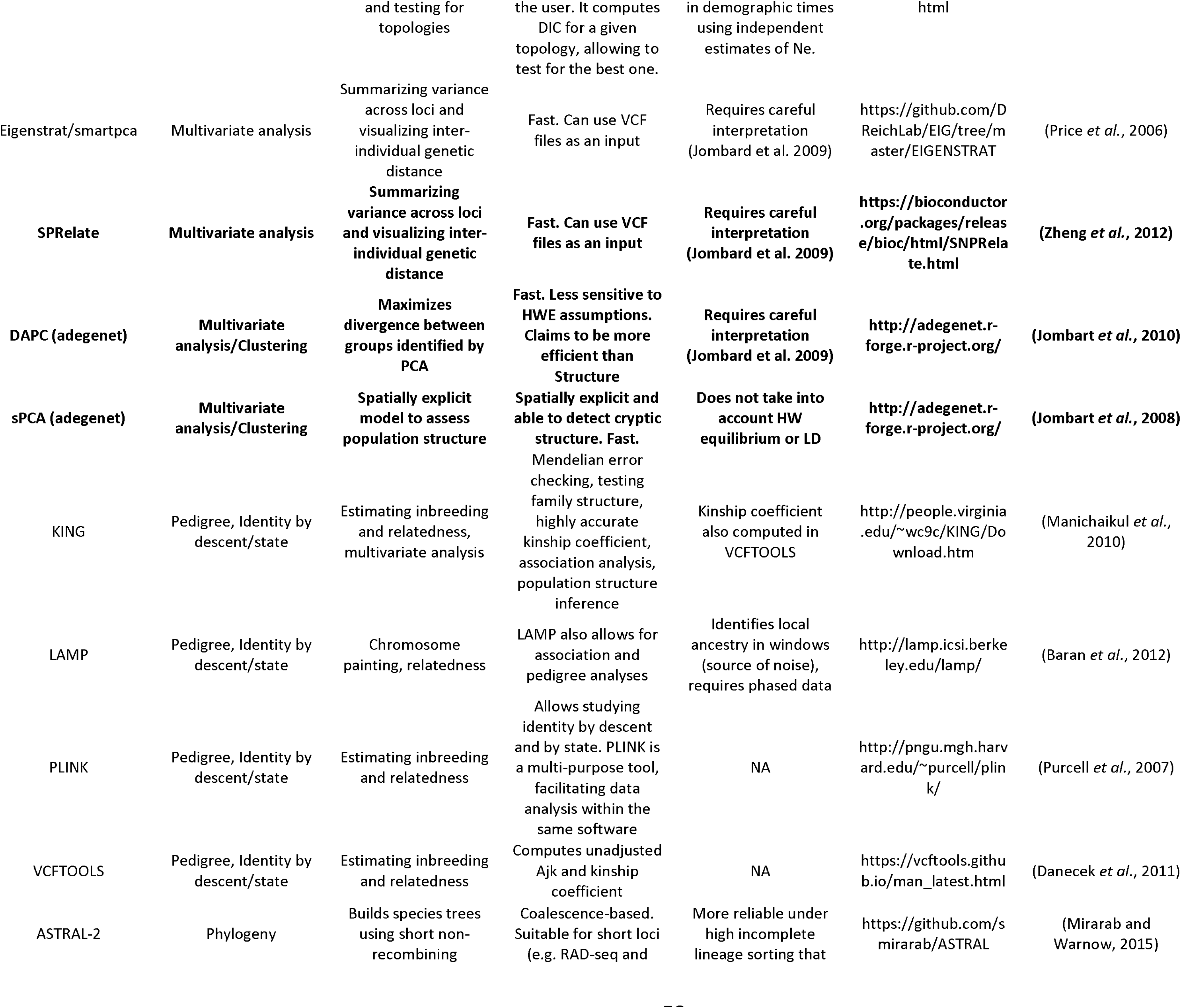
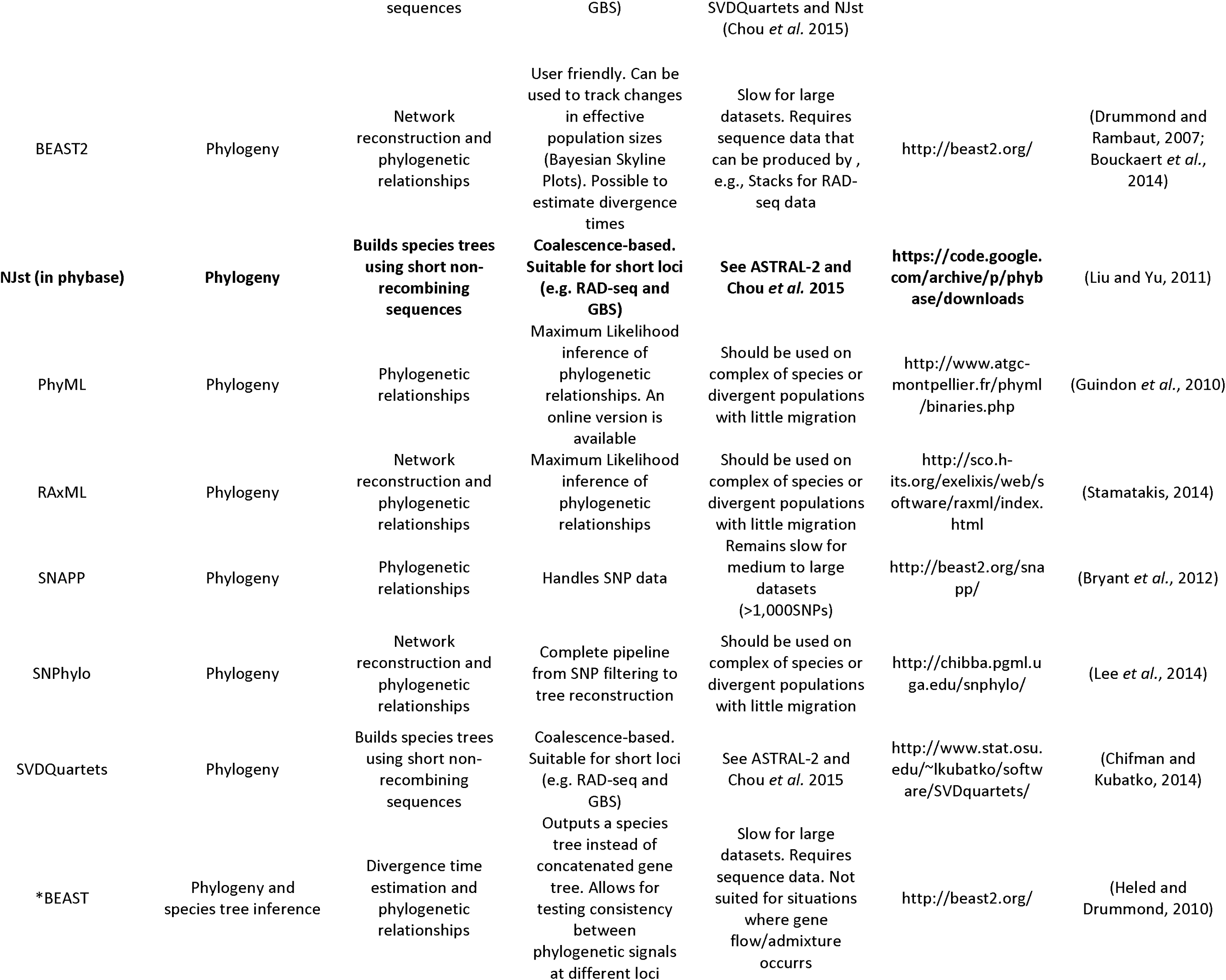
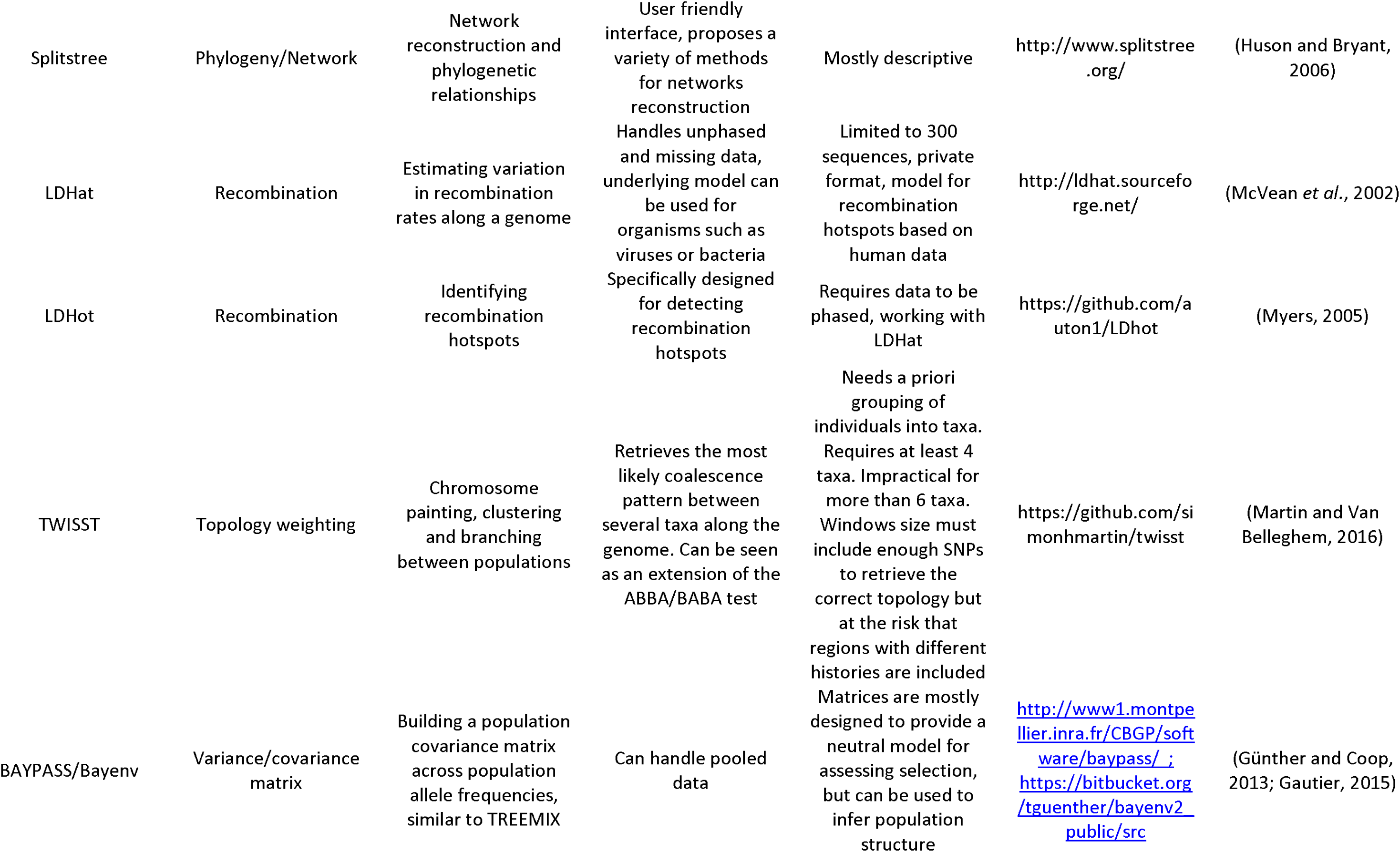
Summary of methods dedicated to data description and assessing population structure. Methods highlighted in bold can be combined in a pipeline within the R software.

In this simplified example, we see that a reciprocal feedback between different aspects of evolutionary genomics is needed (Figure 1). Combining approaches is one of the current grand challenges in evolutionary biology (Cushman, 2014). While large-scale collaborations and sharing of skills between researchers allow for detailed analyses, a regularly updated list of methods would be valuable for smaller research teams to quickly start new projects and evaluate their experimental design.

**Figure 1.**
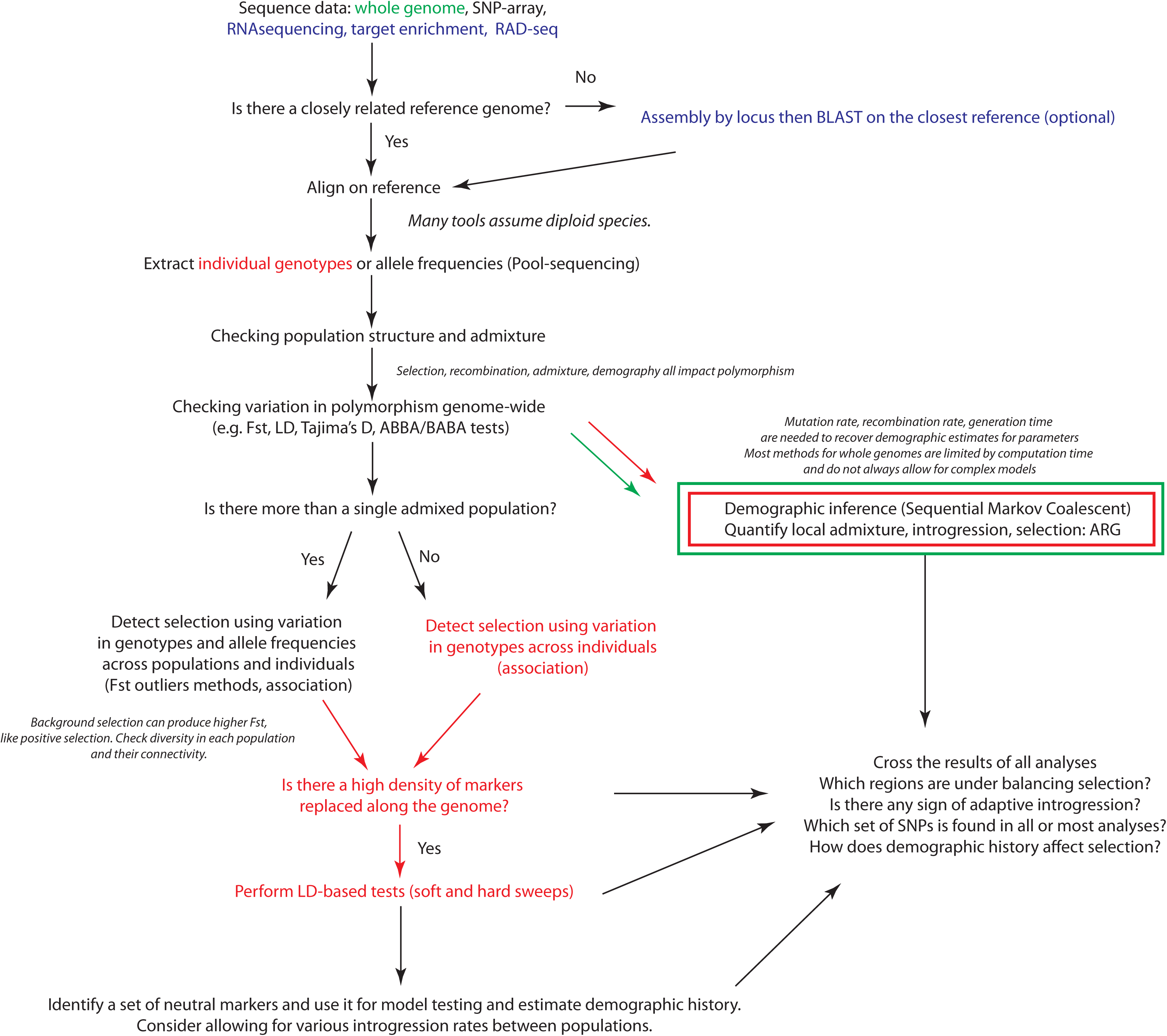
A possible general pipeline for analysing population genomics data using methods described in this paper. In red are indicated options that are generally not suited for pool-seq data. In green are indicated steps that require genome-wide datasets. ARG: Ancestral Recombination Graph (see main text).

In addition to methodological and technical challenges, the widespread use of sophisticated analytical tools is made difficult by the lack of communication between fields (Shafer *et al.*, 2015), little user-friendliness of software, inflation of data formats (Lischer and Excoffier, 2012) and the ever-increasing number of methods made available. Fields like landscape genetics and phylogeography have largely focussed on identifying general patterns in populations history and species diversification. Other researchers are more interested in identifying specific genes that are involved in adaptation in natural populations. All these views contribute to our understanding of causation in biology, an effort that has included genetics, developmental science and ecology (Laland *et al.*, 2011). A global summary of methods used in these different fields would therefore facilitate communication between disciplines.

The last extensive review of methods in population genetics was performed 10 years ago (Excoffier and Heckel, 2006). Since then there has been increasing drive to translate these methods into approaches applicable to genomic data and non-model species. This drive has confirmed the value of population genomics on non-model species in understanding biological diversity at various scales (Mandoli and Olmstead, 2000; Jenner and Wills, 2007; Abzhanov *et al.*, 2008; White *et al.*, 2010; Ellegren *et al.*, 2012; Weber *et al.*, 2013; Poelstra *et al.*, 2014). Such advances are needed to broaden our view about the evolutionary process and improve sampling of distant clades. Ultimately, this process should provide a more balanced picture than the one brought by the study of a few model species (Abzhanov *et al.*, 2008). Genomic approaches also have the potential to improve conservation genetic inference by scaling up the amount of data available (Shafer *et al.*, 2015). Much effort has recently been made in facilitating the diffusion of sometimes complex, state-of-the-art methods. Their application to species with little background data has become more accessible, bringing the potential to add much valuable information.

In this paper, we propose possible pipelines (Figures 1, 2 and 3) to help choose appropriate methods dealing with current questions in population genomics and genetics of adaptation in natural populations. We begin with a succinct review of methods available to obtain genome-wide polymorphism data (Box 1) before focusing on i) methods devoted to the study of population structure and quantitative characterization of population history (Table 1 and 2) and ii) methods aimed at identifying selected loci (Table 3). We end this review by detailing how these analyses can be combined, and present future directions that may be taken by the field of population genomics. The tables and a summary of the methods discussed in this paper will be kept updated to follow improvements, and are available at www.methodspopgen.com.

**Figure 2.**
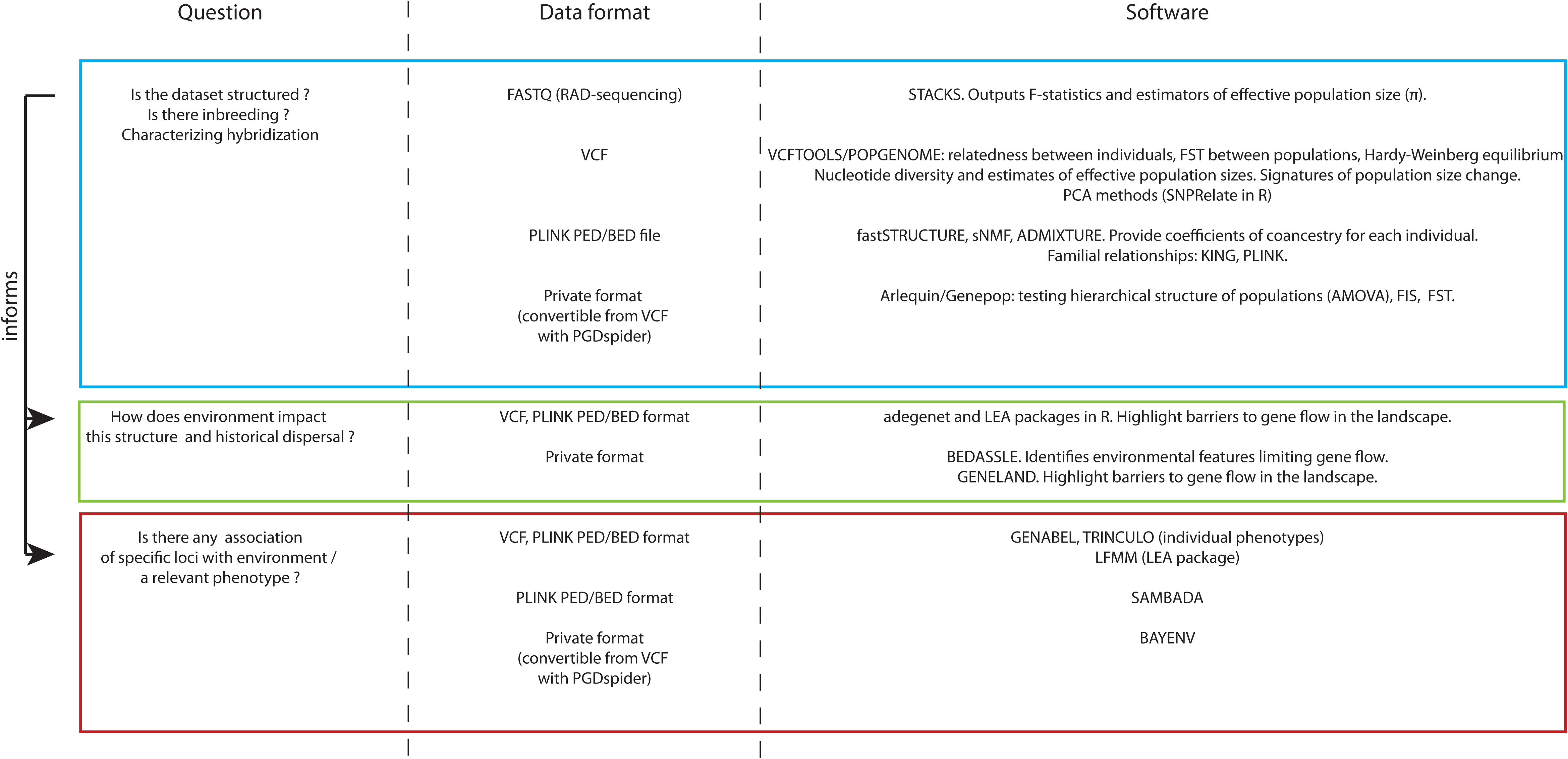
Set of questions and relevant methods to characterize population structure and local adaptation. Proposed methods mostly use common data formats for input files, facilitating their integration in a single pipeline. PGDSpider (Lischer and Excoffier, 2012) can be used to automate file conversion for methods requiring private input format. The proposed methods are not exhaustive, see tables for a more detailed list.

**Figure 3:**
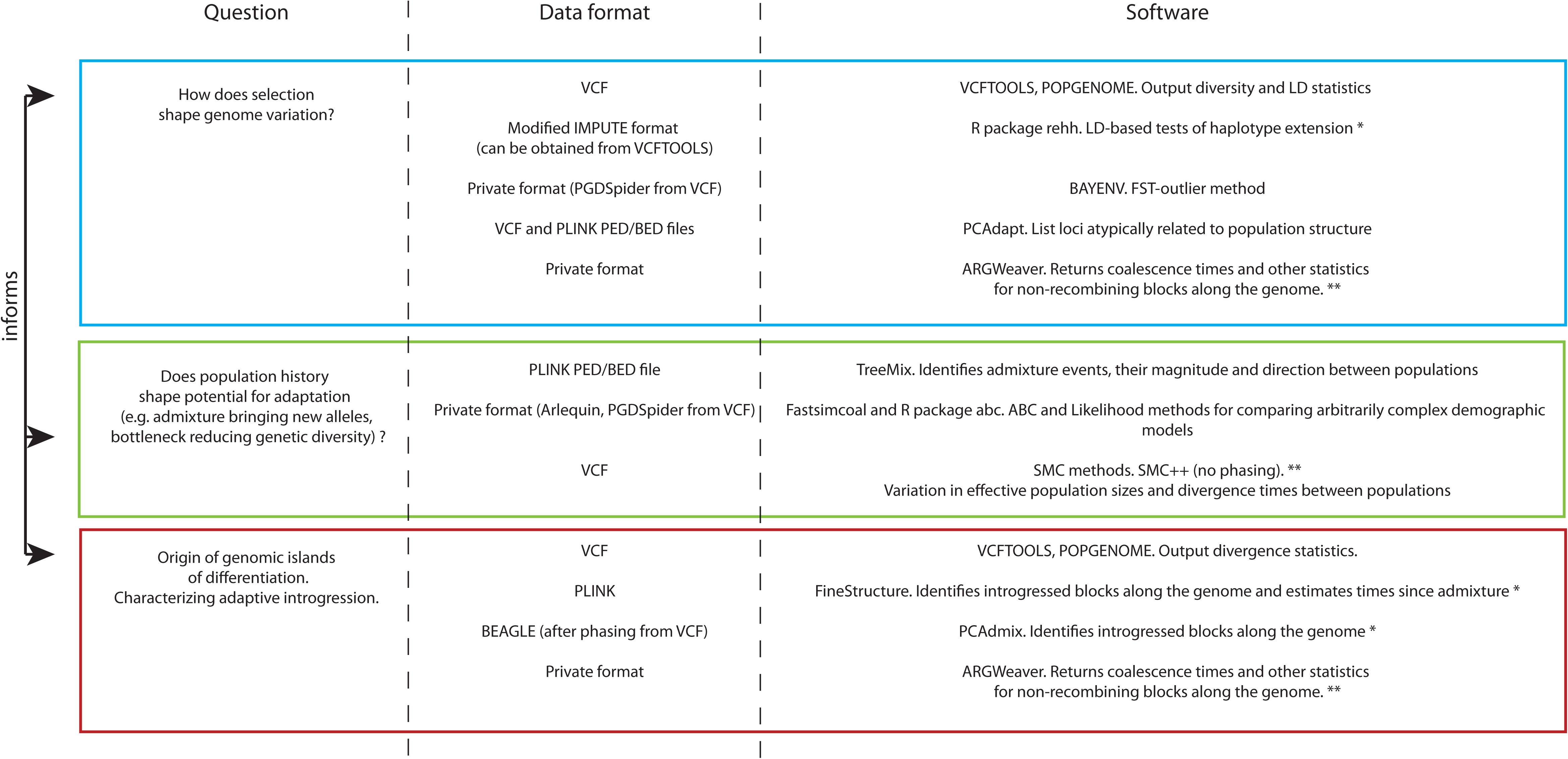
Set of questions and relevant methods to characterize demography and selection. *: requires reference genome; ** requires reference genome and whole genome resequencing.

**Table 2.**
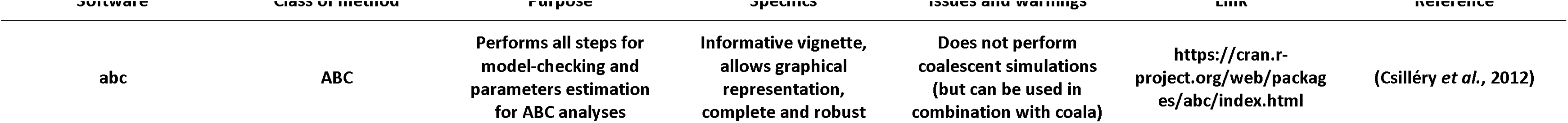
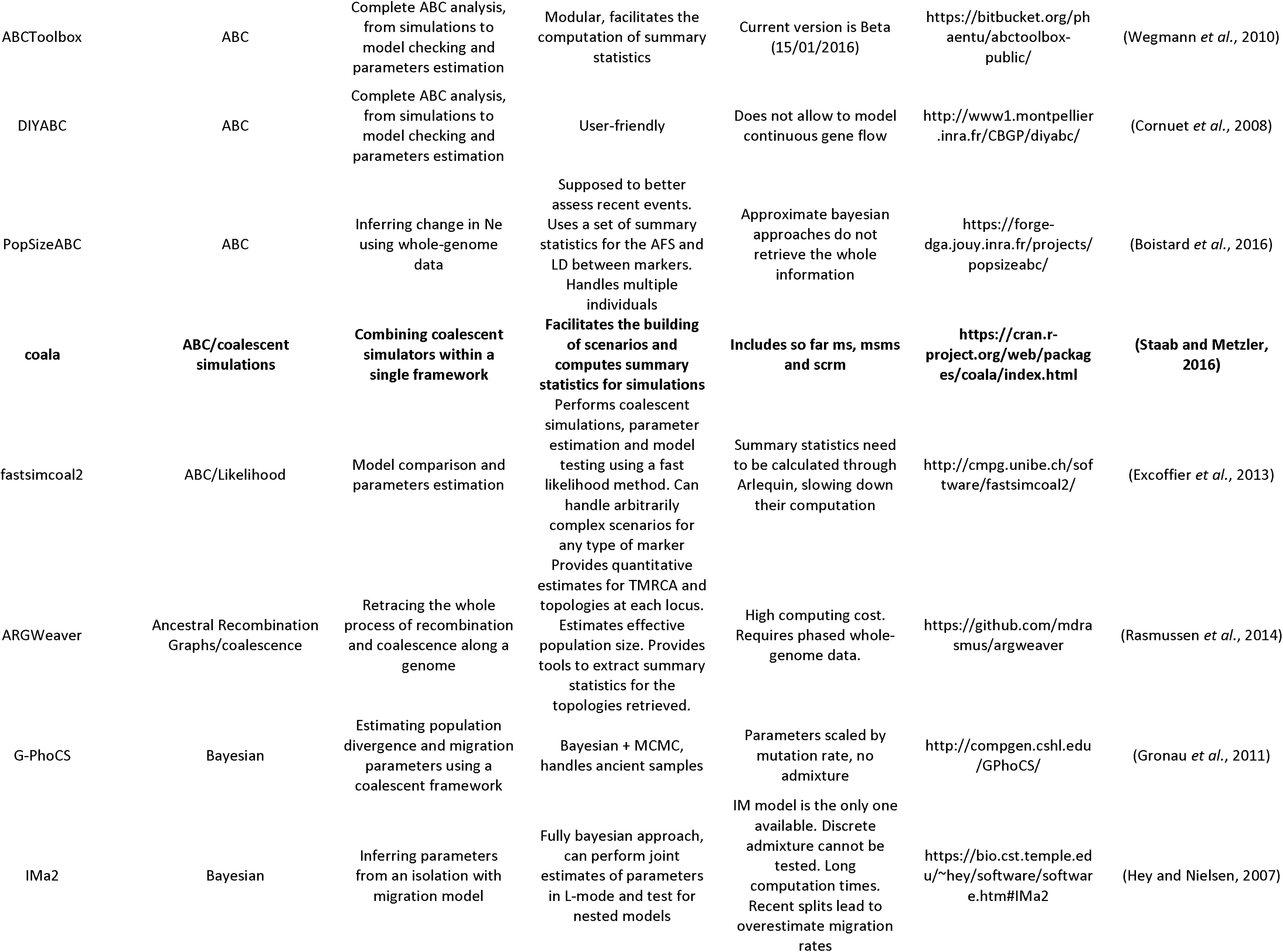
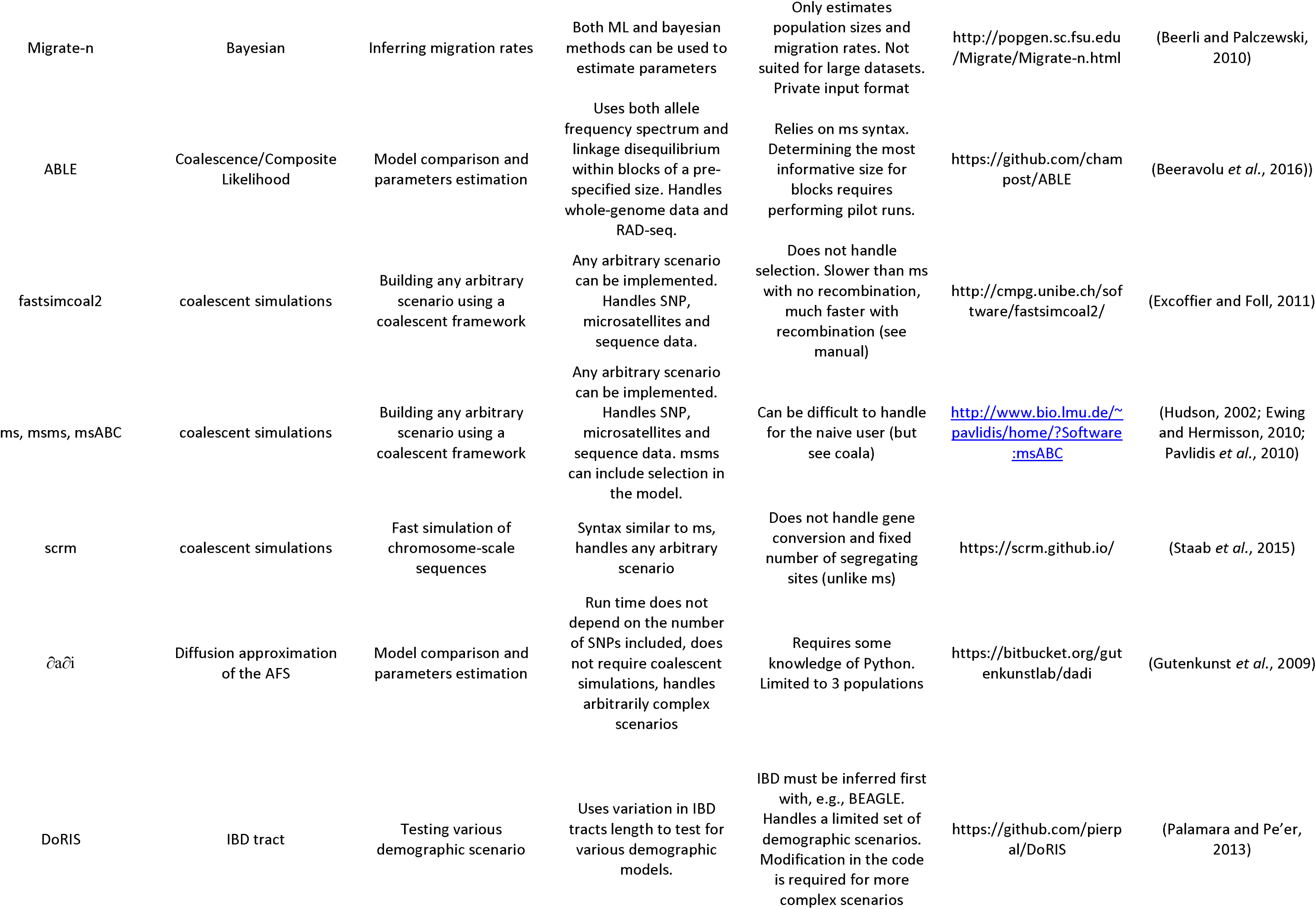
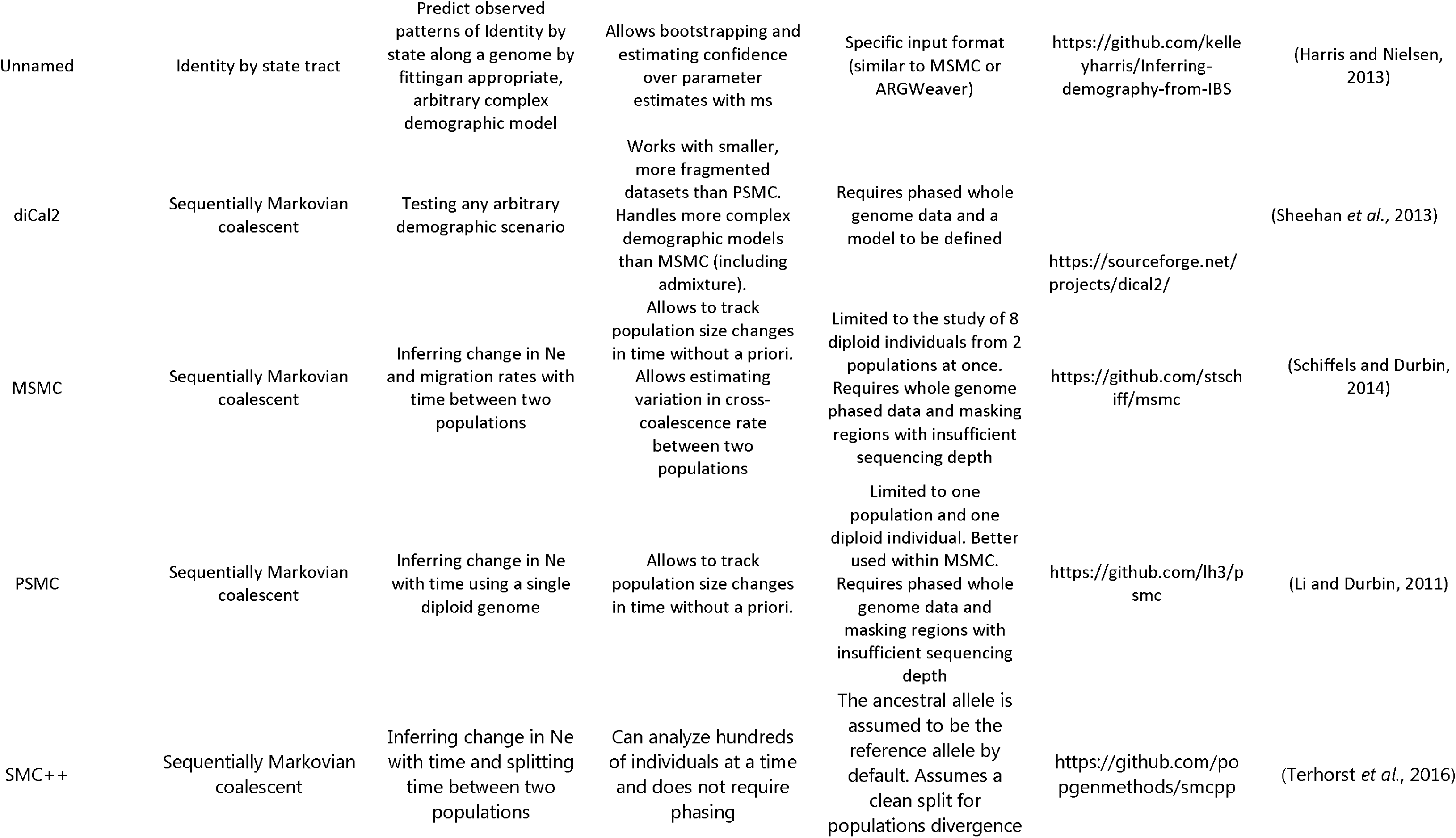
Summary of methods for demographic inference, simulations and scenarios comparisons. Methods available in R are highlighted in bold.

**Table 3.**
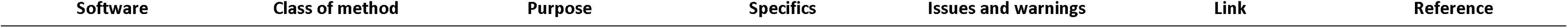
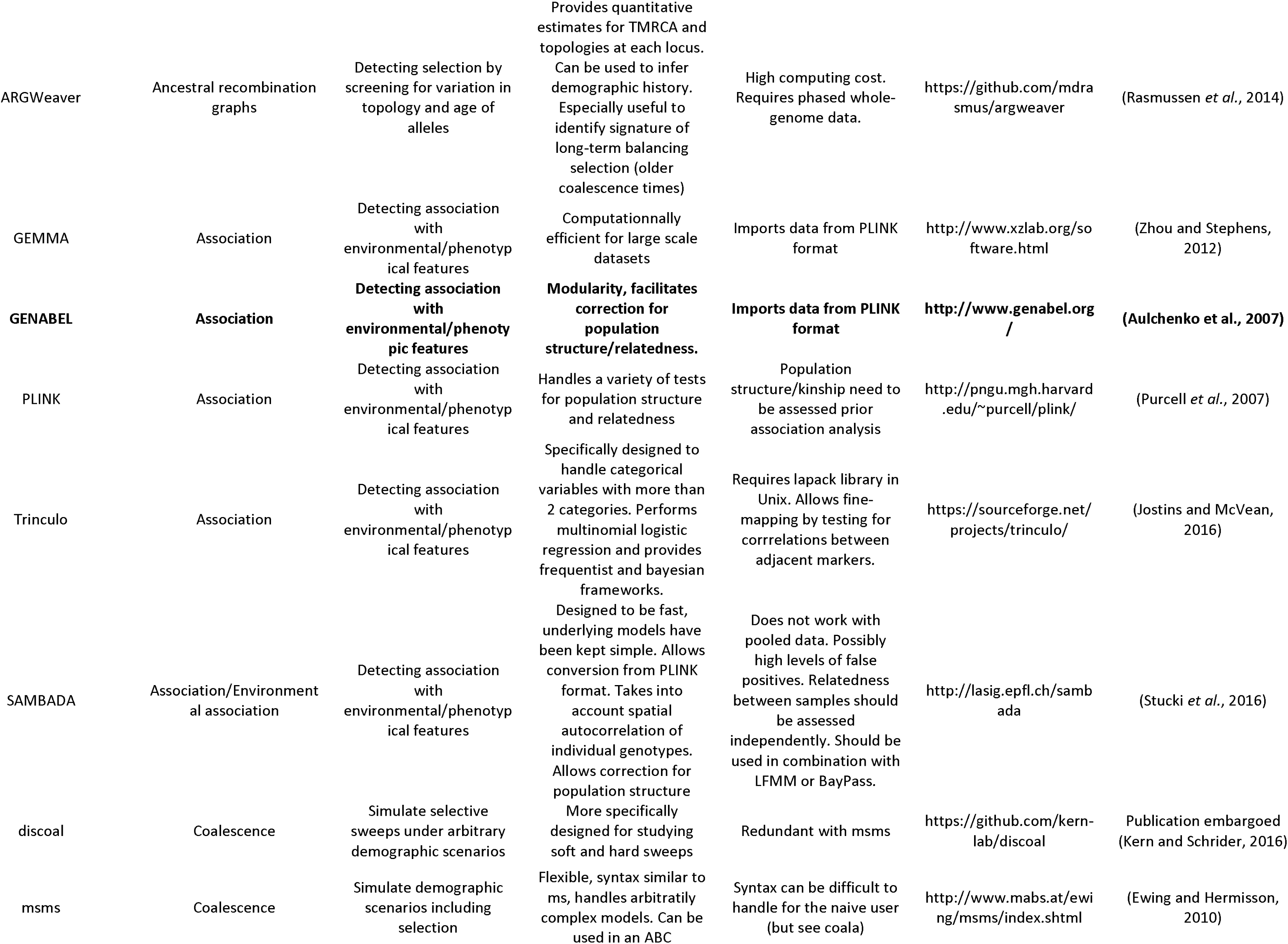
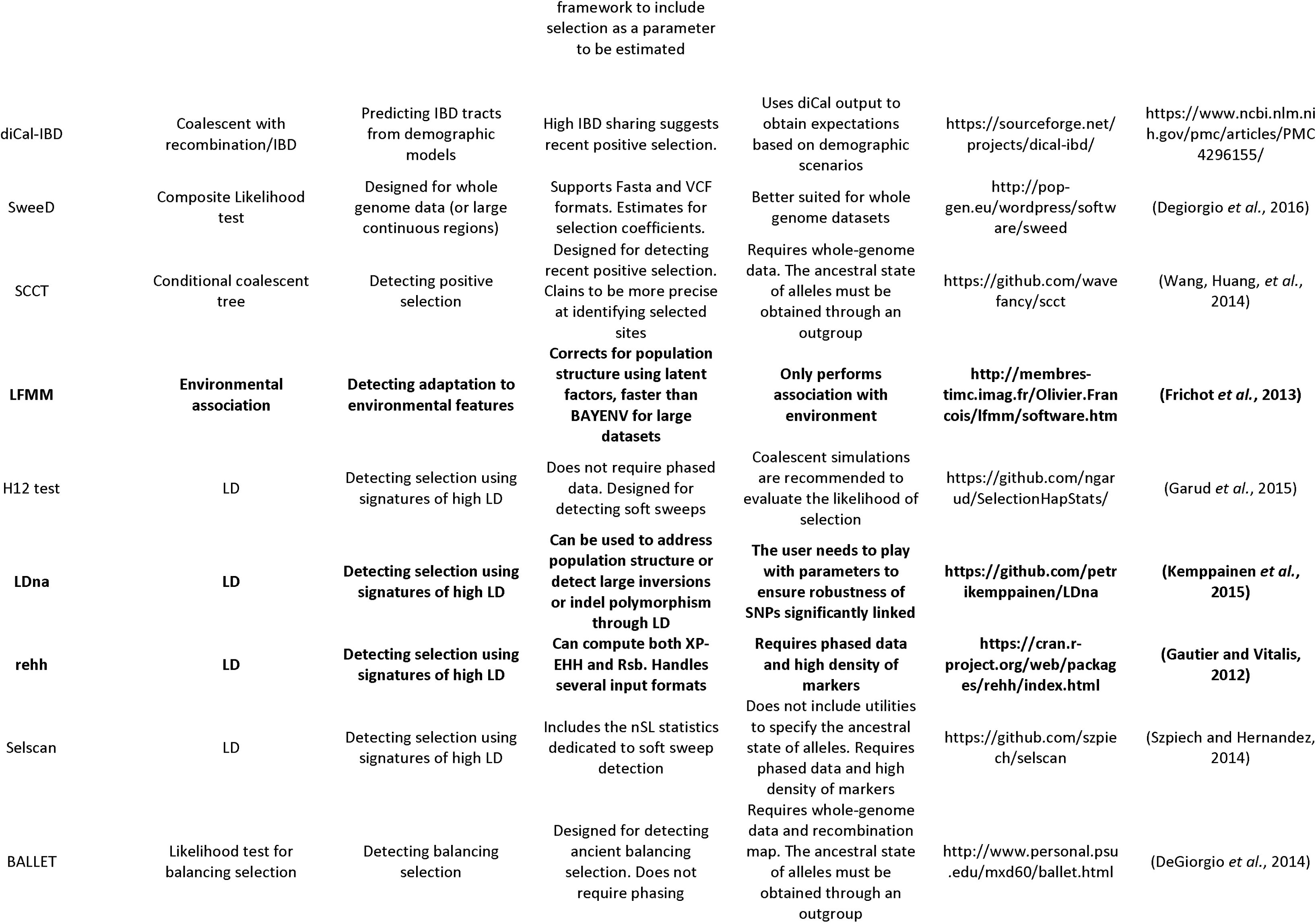
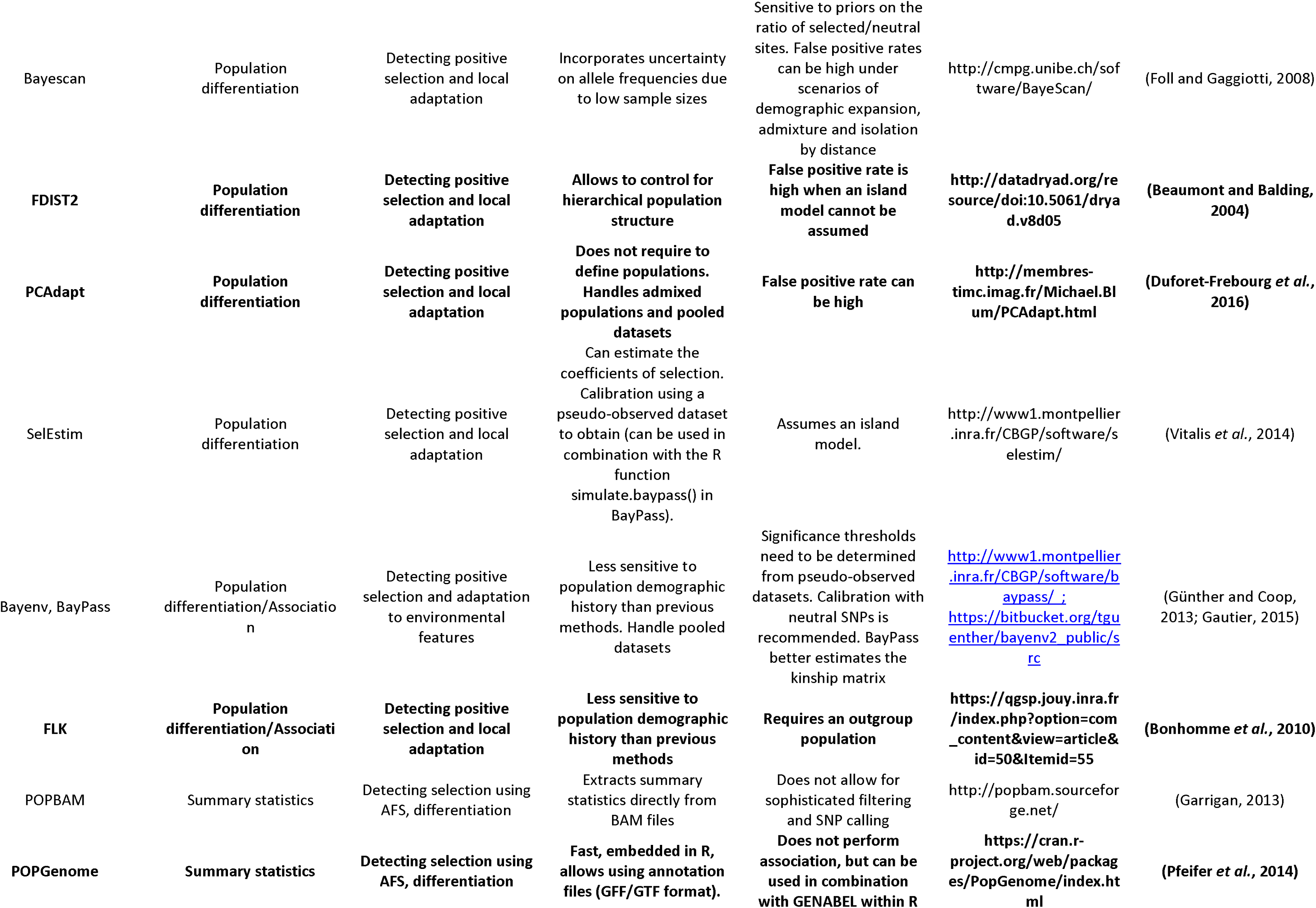
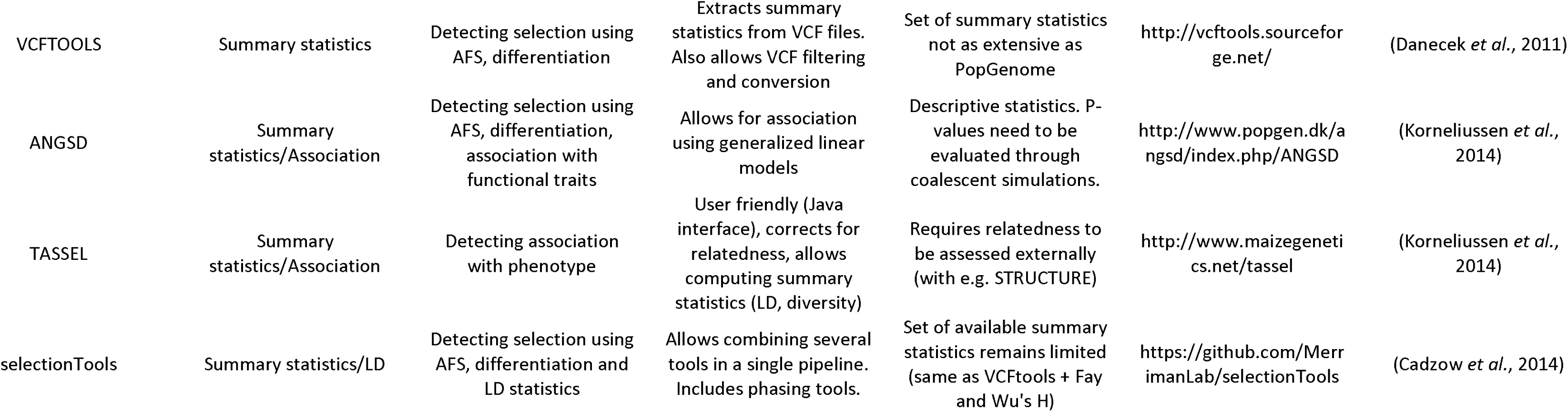
Summary of common methods for identifying loci under selection. Methods available in R are highlighted in bold.

#### Box 1 Common sequencing methods

##### RAD-seq

Reduced representation allows broad sampling of variants across the genome by sequencing DNA fragments flanking restriction sites. Such sampling is not specific to any particular kind of region (e.g. coding or non-coding). Some of the best-known reduced representation techniques include RAD-sequencing (Baird *et al.*, 2008) and Genotyping by Sequencing (GBS; Elshire *et al.*, 2011). Their main interest is their low cost and that they do not require any reference genome (see Davey *et al.*, 2011 for details), although a reference can be useful to identify outlier genomic regions and retrieve linkage disequilibrium information between markers. Use of a reference genome also limits the bias due to paralogy and mapping errors (Hand *et al.*, 2015). Reduced representation allows many individuals to be genotyped at once, and so is widely used for the study of population structure, demography and selection. It does not cover all mutations in the genome and the choice of the restriction enzyme is crucial to control for the density of markers. This choice further controls the mean sequencing depth, the number of mutations close to genes under selection, and the accurate calling of genotypes. The number of SNPs ranges from thousands to millions, which is usually enough to retrieve substantial information about demography and sometimes selection (see Puritz *et al.*, 2014 for a detailed summary of reduced-representation techniques). As a general word of caution, note that RAD-sequencing and related methods display specific properties that can bias genome-wide estimates of diversity, e.g. allelic dropout (Arnold *et al.*, 2013, Puritz *et al.* 2014). However, this type of marker remains valuable for phylogenetic estimation, even for distantly related species (Cariou *et al.*, 2013), and allelic dropout can be compensated for by focusing only on markers sequenced in all individuals. Variations on the original RADseq protocol have been developed to overcome some of these caveats (ddRAD, Peterson *et al.*, 2012; ezRAD, Toonen *et al.*, 2013; 2b-RAD, Wang *et al.*, 2012). Many pipelines have been specifically designed to account for RAD-seq specificities, including Stacks (Catchen *et al.*, 2011), TASSEL-UNEAK (Lu *et al.*, 2013) or TASSEL-GBS for GBS data (Glaubitz *et al.*, 2014).

##### Targeted sequencing

This class of methods allows sequencing and genotyping the same set of genomic fragments or single nucleotide polymorphisms (SNP arrays) across individuals, and has been recently promoted to study non-model species (Jones and Good, 2016). Since the specificity of the probe does not have to be very high, the same probe can be used among closely related species (Nicholls *et al.*, 2015). Conservation of the target genomic region under study is important. High conservation may lead to higher efficiency of capture but can artificially reduce representation of polymorphic regions. Different technologies allow for targeted sequence capture that can be classified by enrichment methods (hybridization-based; PCR-based; molecular inversion probe-based; see Mamanova *et al.*, 2010). Commercial products, such as Agilent’s SureSelect, MYcroarray’s MYbaits or Roche NimbleGen’s SeqCap offer these methods or a derivation (Grover *et al.*, 2012).

Targeted sequencing reduces the genomic representation compared to whole genome sequencing and it allows for multiple individuals to be multiplexed, lowering the cost of sequencing per sample. In addition, the complexity of analysis is reduced compared to whole genome sequencing (WGS), since only a subset of genomic regions is sequenced. By allowing an improvement in spatial and temporal sampling, targeted sequencing can reconstruct dispersal routes and migration between varieties and subspecies (Nadeau *et al.*, 2012; da Fonseca *et al.*, 2016). Another commonly used technique includes Single nucleotide polymorphism (SNP) genotyping arrays have frequently been used in studies aimed at detecting phenotype/genotype associations or to study population struc ture (Gautier *et al.*, 2010; Johnston *et al.*, 2011). However, regenotyping of ascertained SNPs in a new population can lead to bias which can be problematic for demographic inference (Albrechtsen *et al.*, 2010; Lachance and Tishkoff, 2013).

##### RNAseq

RNAseq can be used with and without a reference genome. In the latter case, like any other reduced representation method, it does not provide information of linkage among genes. It has applications on many different evolutionary time scales. Since it mostly sequences coding regions, a deep phylogeny can be constructed with conserved orthologs. Depth of coverage is gene expression dependent, so calling genotypes varies across genes and which must be taken into consideration (Gayral *et al.*, 2013). If a reference genome is available, it is possible to call variants (Piskol *et al.*, 2013). This method is cost-effective and an alternative to whole genome sequencing. However, common variant callers do not behave well with RNAseq due to reads encompassing intronic regions as well as bias introduced during the sequencing library preparation. One of the common variant calling pipelines available is GATK which suggests best practices for calling variants on RNAseq (https://software.broadinstitute.org/gatk/best-practices/). Another variant calling protocol specifically designed for RNAseq is Opossum (Oikkonen and Lise, 2017), which can be used with haplotype-based callers such as Platypus and GATK haplotypeCaller. This software maintains precision and improves the sensitivity of SNP calling compared to the GATK best practice pipeline. RVboost (Wang, Davila, *et al.*, 2014) was developed using the method of variant prioritization, using a so-called boosting method that uses a set of high-confidence variants to set a model of good quality variants. All RNA variants are then prioritized and called based on this model. It outperforms Variant Quality Score Recalibration (VQSR) from the Genome Analysis Tool Kit (GATK) and the RNA-Seq variant calling pipeline SNPiR (Piskol *et al.*, 2013). RVboost can indentify false variants introduced by random hexamer priming during library preparation.

##### Whole genome resequencing

Whole-genome resequencing requires a well assembled reference and is more expensive than RAD-seq or targeted sequencing, especially for species with long and complex genomes. Some methods do not actually require any reference sequence to call SNPs from raw reads, like kSNP2 (Gardner and Hall, 2013) or DiscoSNP (Uricaru *et al.*, 2015). However, this limits the main interest of this approach, since mapping back on a reference has the potential to provide a complete overview of structural and coding variation. It also allows the use of powerful methods to track signatures of selection (see below). Pooled sequencing (Futschik and Schlötterer, 2010) can be an option to reduce costs, but generally restricts analyses to methods focusing on allele frequencies. Since individual information is not available, variation in Linkage Disequilibrium across individuals (LD) cannot be exploited. Shallow sequencing (1-5X per individual) may be a way to partly overpass this last issue for a similar cost (Buerkle and Gompert, 2013), but should not be used for methods requiring phasing and unbiased individual genotypes.

Shallow shotgun sequencing also allows retrieving complete plastomes, due to the representation bias of mitochondrial or chloroplast sequences. Plastome sequences can provide insightful information into the evolutionary history of populations or species, and recent work has successfully used shallow sequencing to reconstruct mitochondrial or chloroplast sequences in plants (Malé *et al.*, 2014), animals (Hahn *et al.*, 2013) or old and altered museum samples (Besnard *et al.*, 2016). Methods such as MITObim (Hahn *et al.*, 2013) provide an automated and relatively user-friendly way to reconstitute plastome sequences, which can then be analyzed as a single non-recombining marker for phylogeny or population genetics.

## Population structure and data description

### Population structure and diversity

Description of the data is essential to assess the proportion of loci displaying a consistent pattern, and characterize how genetic diversity is partitioned within species. Genetic diversity and its genome-wide variance are directly impacted by variation in many factors including effective population sizes, population structure, inbreeding, migration, and recombination rates. Their characterization must be performed prior to any analysis to get insights into the forces and constraints acting on populations.

A key aspect when describing a new dataset is the assessment of relatedness between individuals or localities. Neglecting population structure can dramatically bias demographic inference, especially when gene flow is not accounted for or panmixia is assumed (Chikhi *et al.*, 2010; Heller *et al.*, 2013). It also biases the detection of loci under selection (e.g. Nielsen *et al.*, 2007). Cryptic population structure is typically a confounding effect in studies of phenotype-genotype association studies, when a given feature or trait is disproportionally found in a population or a set of related individuals (Balding, 2006). Fortunately, the abundance of SNP data produced by typical genomic studies is often enough to thoroughly assess relatedness between individuals.

Many tools currently exist to infer population structure (Table 1, Figure 2). An elegant and efficient class of methods relies on using multivariate approaches such as principal component analysis (PCA) to infer relatedness between individuals and populations without *a priori* knowledge. Since these methods do not have underlying assumptions based on population genetics, they are suitable for analyzing species displaying polyploidy or mixed-ploidy (Dufresne *et al.*, 2014). A detailed review of these methods has been already performed (Jombart *et al.*, 2009) and an exhaustive list of their applications is beyond the scope of this review. These approaches have been especially useful to study the consistency between geographical and genetic structure in human populations of Europe (Novembre *et al.*, 2008). They were also recently applied to RAD-sequenced populations of a freshwater crustacean (*Daphnia magna*). Procrustes rotation (Novembre *et al.*, 2008) was used to match geographical coordinates with PCA axes, showing how isolation by distance has shaped genetic structure (Fields et al. 2015).

Methods for estimating the relatedness of individuals are suited to studies relying on pedigree information, or if there are reasons to suspect that familial relationships can play a major role in shaping genetic structure of the population(s) considered. When each individual in a study is sampled from a different location or environment, estimating relatedness also provides a way to assess the genetic distance between individuals. Genetic distance can then be compared with geographical or ecological distance. For example, in a recent study using more than 1000 *Arabidopsis thaliana* genomes, estimates of relatedness have allowed the identification of putatively relictual populations that may have persisted in Europe since the last Ice Age (Alonso-Blanco *et al.*, 2016).

Approaches such as Structure (Pritchard *et al.*, 2000) and fastSTRUCTURE (Raj *et al.*, 2014) have been widely used to determine hierarchical population structure and admixed populations by grouping individuals in clusters. The optimal number of clusters (K) can then be determined based on likelihood, although examining population structure for a range of K can allow substructure to be better identified. The main interest of these approaches is that they provide a measure of coancestry coefficients, which are the proportions of an individual genome originating from multiple ancestral gene pools. Such information is more difficult to retrieve with approaches such as PCA. There have been criticisms however about whether ambiguous assignment could be actually interpreted as a signal of admixture, and detailed inference requires thorough model testing and estimating the goodness of fit of a model with admixture (see Falush *et al.*, 2016).

### Heterogenous patterns of divergence between species along their genomes

Advantageous alleles can migrate from one population to another, resist introgression from other populations, reach fixation and erase diversity around them. This is one scenario leading to heterogenous patterns of divergence along the genome, the so-called islands of divergence (Wolf and Ellegren, 2016). Alternative scenarios leading to similar patterns were recently highlighted (Cruickshank and Hahn, 2014). Understanding the origin of genomic regions under selection highlights the evolutionary history of adaptive alleles (e.g. Abi-Rached *et al.*, 2011) and contributes to our understanding of the origin and maintenance of reproductive isolation. Studies focusing on hybrid zones and introgression have provided inspiring examples (Hedrick, 2013), as demonstrated by recent work focusing on patterns of heterogenous gene flow in *Mytilus* mussels (Roux *et al.*, 2014), localized introgression and inversions at a color locus in *Heliconius* butterflies (The Heliconius Genome Consortium *et al.*, 2012) and adaptive introgression of anticoagulant resistance alleles in mice (Song *et al.*, 2011). Descriptive statistics computed along genomes provide valuable information in this context. One may for example plot the distribution of a differentiation measure such as FST (Weir and Cockerham, 1984) between populations, mean linkage disequilibrium or nucleotide diversity. Such an approach has been used in *Ficedula* flycatchers, which uncovered clear genomic islands of divergence and the higher differentiation on sexual chromosomes due to ongoing reproductive isolation (Ellegren *et al.*, 2012). Other approaches, such as chromosome painting (Table 1), extend PCA and Structure-like methods by incorporating information about the relative order of markers in the genome, allowing identification of regions for which ancestry differs from the rest of the genome.

### Heterogeneous structure in space: landscape genomics

Landscape (as well as seascape and lakescape) genetics has widely contributed to our understanding of how ecological and geographical variation affects species history and adaptation (Manel and Holderegger, 2013). Of central importance in this field is the identification of how populations are connected and how organisms move in the landscape matrix. Environmental heterogeneity has a strong impact on how genetic diversity is shaped by migration success between populations, for example after a range expansion (Wegmann *et al.*, 2006). A spatially explicit perspective provides context to understand the evolution of locally adapted genes. Moreover, identifying how and where populations (or closely related species, see Roux et al. 2016) hybridize is crucial when it comes to characterizing colonization trajectories, tension zones and secondary contacts (Gay *et al.*, 2008; Bierne *et al.*, 2011).

Some methods can explicitly use spatial information to inform clustering, allowing improved consideration of the effect of landscape heterogeneity on selection against migrants and drift. This spatial perspective can be useful to visualize the location and shape of hybrid zones (Guedj and Guillot, 2011). Landscape genetics has valuable application in management and conservation, where it is useful to identify the relevant evolutionary significant units displaying spatial and ecological divergence. Furthermore, researchers are often interested in testing the impact of ecological variation on genetic structure. Mantel tests have been popular to investigate relationships between ecological variables and genetic differentiation while accounting for geographical distances. However, these tests are biased by spatial autocorrelation, assume linear dependence between variables, and do not allow testing the relative contribution of each variable (Legendre and Fortin, 2010; Guillot and Rousset, 2013). Methods such as BEDASSLE (Bradburd *et al.*, 2013) can be used to complement these approaches, and identify which combination of geographical and ecological distance limits dispersal. However, disentangling these effects has proved to be complex and a deeper analysis of genes more strongly impacted by either geography or ecology may be more informative when it comes to the proximate causes of reduced dispersion and differentiation, such as biased dispersal (Edelaar and Bolnick, 2012; Bolnick and Otto, 2013) or selection against migrants (Hendry, 2004). Landscape genomics now extends its focus to adaptive genetic variation, and benefits from new methods targeting signatures of selection (Figure 2 and below).

## Population history

### Phylogeny

Phylogenetics has a long history that is linked to the broader topic of systematics (Moritz & Hillis 1996; Baum & Smith 2013). Since their inception in the 1980s, molecular phylogenetic methods have been used to address a wide range of problems at different taxonomic scales, including intraspecific population history. Recent advances in molecular phylogenetic methods, and the employment of different types of NGS data is well beyond the scope of this review (see e.g. Moriarty Lemmon & Lemmon 2013; Cruaud et al. 2014; Wen et al. 2015). Rather we focus on the use of phylogeny within the context of studies of intra-specific population history and selection. In this respect, both Maximum Likelihood and Bayesian approaches have become popular to investigate evolutionary relationships between individuals from different populations, even when divergence is very recent (e.g. Wagner *et al.*, 2013). These methods are implemented in softwares such as RAxML (Stamatakis, 2014) and BEAST2 (Drummond and Rambaut, 2007). Ultimately, all molecular phylogenies reconstruct the geneaology of the genes with which they have been constructed. Therefore, a basic assumption when using them to infer lineage history at any taxonomic level (populations, species, and higher taxonomic units) is that the gene tree is representative of lineage history. This assumption is likely to be particularly weak at the population level, since the influences of gene flow, selection, and incomplete lineage sorting are strong at this scale, and may cause gene trees to deviate from population history. Nonetheless, such phylogenies can provide a useful starting point for inferences that are complemented with other methods.

When using genome-wide data at the population level, methods specifically dedicated to reconstructing multiple species coalescent models (MSC) such as *BEAST (STAR-BEAST) should be preferred over concatenation (Edwards *et al.* 2016), since they allow discordance between species trees and individual gene trees to be identified. Note that these methods can be strongly biased when it comes to estimate divergence times and effective population sizes (Leaché *et al.*, 2014). The impact of gene flow and recombination on phylogenetic methods is however an alley of research that will allow better integration between phylogeny and population genetics (Edwards *et al.*, 2016). Such integration is particularly needed for species and populations that are in the “grey zone of speciation” (Roux *et al.*, 2016). Recent advances in MSC methods handling extremely short, non-recombining fragments (see Chou *et al.*, 2015 for a comparison) are promising, especially for datasets such as those produced by GBS.

While useful to infer topologies, caution is advised when using branches lengths obtained from SNP-only datasets, e.g. to calculate divergence times between different groups or species (Leaché *et al.*, 2015). For this purpose, it might therefore be easier to extract from the data both variant and invariant sites at several genes or RAD contigs, and analyze the whole sequences in a software like BEAST2. Network methods implemented in Splitstree (Huson and Bryant, 2006), make less assumptions and account for potentially conflicting signals due to high gene flow. Unfortunately, such methods remain mostly descriptive.

### Approximate Bayesian Computation

Phylogenetic methods tend to be slow for large datasets, and generally do not attempt to account for many effects that are crucial in population genetic interpretation, such as gene flow and recent demographic events within species. A more suitable framework for microevolutionary studies relies on coalescence theory. Population geneticists first developed coalescent theory as a way of modeling the genealogy of alleles from a sample of a large population. Going backward in time, alleles merge (coalesce) in a stochastic way until reaching their most recent common ancestor (Kingman, 1982). Obtaining demographic estimates (e.g. time in years) for parameters usually requires that mutation rate and generation time be known or at least reasonably well estimated, for example from closely-related species with similar life history.

Computationally fast approaches include Approximate Bayesian Computation (ABC), which compares the empirical data with a set of simulated data produced by coalescent simulations under scenarios predefined by the user (Table 2). By measuring the distance between carefully chosen summary statistics describing each simulation with those from the observed dataset, it is possible to infer which scenario explains the data the best. More information on how to perform ABC analyses are described by Csilléry *et al.* (2010). The main advantage of ABC is that it allows handling any type of marker and arbitrarily complex models, contrary to methods like IMa where the model is predefined. However, using summary statistics leads to the loss of potentially useful information (Robert *et al.*, 2011).

### Likelihood methods based on the allele frequency spectrum (AFS)

Recently, new likelihood methods based on the AFS emerged to facilitate and speed up the analysis of large SNP datasets. Different patterns of gene flow and demographic events all shape the AFS in specific ways (e.g. alleles are likely to occur at more similar frequencies if divergence is recent or if populations are highly connected). These approaches quickly estimate parameters using composite likelihoods, and do not explicitly take into account correlations induced by LD between physically linked markers (but see ABLE, Table 2). This might limit power to detect recent demographic events (e.g. migration, Jenkins *et al.*, 2012). Including SNPs that are physically close together should not strongly bias parameter estimation. However, such an approach prevents direct comparisons of likelihoods from different models. Therefore, physically independent SNPs should be used to consider composite likelihoods as quasi likelihoods for model comparison (Excoffier *et al.*, 2013). Note that the AFS can also be used as a set of summary statistics for ABC inference. Using allele frequencies estimated from pooled datasets is also feasible, as illustrated by a recent study on hybridization in *Populus* species where AFS was estimated from pooled whole genome resequencing data (Christe *et al.*, 2016).

The number of mutations found in a given length of DNA sequence directly depends on the mutation rate. One drawback when using SNP data without considering monomorphic sites is that the mutation rate per generation can not be used to convert parameters into demographic estimates (Excoffier *et al.,* 2013). Another possibility consists of calibrating parameter estimates by including a fixed parameter in the analysis, such as population size or divergence time. An issue specific to SNP arrays is ascertainment bias, which is the systematic deviation of allele frequencies from theoretical expectations due to the choice of individuals used at the step of SNP discovery. For example, if SNPs found in one population are the only ones genotyped in another population, a whole set of markers polymorphic in the second population but not in the first will be missed, biasing the AFS (Lachance and Tishkoff, 2013).

Reaching a high level of precision when estimating demographic parameters can be challenging when information is lacking about the evolutionary history of the species considered. However, even when such information is lacking it is possible to compare the likelihoods of different demographic scenarios, a procedure that has been successfully applied to many species to shed light on the process of speciation (Roux *et al.*, 2016).

### Methods using whole-genome resequencing

Recently, methods have been developed to infer variation in population sizes with time using the whole genome of just one diploid individual. This began with the Pairwise Sequentially Markovian Coalescent (PSMC, Li and Durbin, 2011), and extensions have been made to this model to allow for several genomes. Such methods have the advantage of requiring only a few individuals, and no *a priori* knowledge of population history. One general drawback, however, is that they are limited to rather simple scenarios, and do not handle more than two populations as yet (but see diCal2, Table 2). While powerful, they are sensitive to confounding factors such as population structure (Orozco-terWengel, 2016) that lead to false signatures of expansion or bottleneck. They also do not allow extremely recent demographic events to be investigated, since the coalescence of two alleles from a single individual in the recent past (a few tens to hundreds generations) is infrequent. Moreover, most of these methods require the data to be phased (but see SMC++, Table 2), for example with fastPhase (Scheet and Stephens, 2006) or BEAGLE (Browning and Browning, 2011). In addition, phasing errors can lead to strong biases in parameters estimates for recent times (Terhorst *et al.*, 2016). An extension of these methods takes into account population structure and aims to identify the number of islands contributing to a single genome, assuming it is sampled from a Wright n-island meta-population (Mazet *et al.*, 2015). Such developments should improve the amount of information retrieved from only a few genomes. However, natural populations are structured and connected in complex ways, which can bias demographic inferences, even for popular markers such as mitochondrial sequences (Heller *et al.*, 2013).

Methods based on tracts of identity-by-descent (IBD, Palamara and Pe’er, 2013) constitute an interesting alternative for more complex model testing when whole genome or densely genotyped datasets are available in large number. Such methods allow recent demographic events to be inferred with relative precision. They are used to predict the length of haplotypes shared by two individuals that are inherited from a common ancestor without recombination. However, IBD detection requires large cohorts and accurate phasing, and therefore application of these methods has been largely restricted to human populations so far (Browning and Browning, 2011; Palamara and Pe’er, 2013). Another approach has used tracts of identity-by-state to perform demographic inference over a range of time-scales (IBS, Harris and Nielsen, 2013). IBS tracts are directly observable since they are simply the intervals between pairwise differences in an alignment of sequences and do not require any assumption about coancestry to be defined. The method predicts the length distribution of IBS tracts for pairs of haplotypes under a range of demographic parameters. These predicted spectra are then compared to empirical data under a likelihood framework, as with methods based on the AFS.

There is currently a tradeoff to be made between methods allowing for arbitrarily complex models that are defined *a priori* by the user (e.g. ABC), and methods that allow population history to be inferred agnostically (e.g. PSMC). While the first category of methods are typically the highest performers at inferring complex population history from a moderate number of markers, it is currently only the second category of methods that are able to make use of the full information provided by whole genome data. Using both methods can therefore help in accurately retrieving the evolutionary history of a given species. For example, a recent study on maize demographic and selective history used both ∂a∂i and and Markovian Coalescent methods to characterize the bottleneck and expansion associated with domestication (Beissinger *et al.*, 2016).

## Screening for selection and association

### Selection and its impact on sequence variation

While demographic forces such as drift and migration will affect the whole genome, selection is expected to be specific to particular portions of the genome, and therefore yield discrepancies with genome-wide polymorphism (Lewontin and Krakauer, 1973). Selection affects allele frequencies and polymorphism in predictable ways at the scale of single populations (Charlesworth, 2006; Charlesworth and Charlesworth, 2010). Several statistics summarize them, such as π, the nucleotide diversity (Nei and Li, 1979), Tajima’s D (Tajima, 1989), and Fay and Wu’s H (Fay and Wu, 2000). Using a combination of these statistics allows targets of selection to be identified with greater precision, and minimizes the confounding effects of demography (Nielsen *et al.*, 2005). This approach has been used to develop composite tests, such as the composite likelihood ratio (CLR) test (Nielsen *et al.*, 2005) that aim to detect recent selective sweeps.

### Methods based on population subdivision

When an allele is under positive selection in a population, its frequency tends to rise to fixation, unless gene flow from other populations or strong drift prevents this from happening (Charlesworth *et al.*, 1997). It is therefore possible to contrast patterns of differentiation between populations adapted to their local environment to detect loci under divergent selection (e.g. displaying a high F_st_). However, it is essential to control for population structure, as it may strongly affect the distribution of differentiation measures and produce high rates of false positives. First attempts to take into account population structure and variation in gene flow included FDIST2 (Beaumont and Nichols, 1996). This method models populations as islands and is aimed at detecting loci under selection by contrasting heterozygosity to F_st_ between populations. More sophisticated methods are now available (Table 3), dedicated to the detection of outliers in large genomic datasets. Most of them correct for relatedness across samples, and can test association between allele frequencies and environmental features (see the extensive review by François *et al.*, 2015). These methods are particularly well suited for the study of RAD-sequencing data, for which allele frequencies are often the only information available in the absence of any reference genome.

Detecting association between environment and allele frequencies does not necessarily imply a role for local adaptation. For example, in the case of secondary contact, intrinsic genetic incompatibilities can lead to the emergence of tension zones that may shift until they reach an environmental barrier where they can be trapped (Bierne *et al.*, 2011). Characterizing population history is required to draw conclusions about the possible involvement of a genomic region in adaptation to environment. The sampling strategy must take into account the particular historical and demographic features of the species investigated to gain power (Nielsen *et al.*, 2007). The sequencing strategy must also be carefully considered to control for spatial autocorrelation of genotypes due to isolation by distance and shared demographic history.

### Genome-wide association

The methods described above focus on allele frequencies at the population scale, but do not test association with traits that vary between individuals within populations (e.g. resistance to a pathogen, symbiotic association, individual size or flowering time). For this task, methods performing Genome-wide association analysis (GWAS) are better suited. The recent development of multivariate methods such as PCAdapt (Duforet-Frebourg *et al.*, 2016) also allow loci putatively under selection to be identified in admixed or continuous populations without requiring information about individual phenotype.

Uncovering the genetic basis of complex, polygenic traits remains challenging, even in model species (Pritchard and Di Rienzo, 2010; Rockman, 2012). It may be unavoidable as a first step to focus only on traits that are under relatively simple genetic determinism. This can, however, lead to the overrepresentation of loci of major phenotypic effect, a fact that should be acknowledged when discussing the impact of selection on genome variation. The fact that loci of major effect are the easiest to target does not imply that they are necessarily the main substrate of selection (Rockman, 2012). Association methods may help targeting variants undergoing soft sweeps, weak selection or those involved in polygenic control of traits (Pritchard *et al.*, 2010). In such cases, signatures of selection may be subtle and sometimes difficult to retrieve from allele frequency data.

### Detecting selection with methods focusing on LD

LD is increased and diversity is decreased near a selected allele, especially after recent selection. A class of methods are aimed at targeting those regions that display an excess of long homozygous haplotypes, such as the extended haplotype homozygosity (EHH) test (Sabeti *et al.*, 2002). It is also possible to compare haplotype extension across populations, with the Cross Population Extended Haplotype Homozygosity test (XP-EHH (McCarroll *et al.*, 2007)) or Rsb (the standardized ratio of EHH at a given SNP site (Tang *et al.*, 2007)). Individuals included in the analysis should be as distantly related as possible to improve precision and avoid an excess of false positives. These methods require data to be phased in order to reconstruct haplotypes. Statistics dedicated to the detection of selection on standing variation or on multiple alleles (so called soft sweeps) are also available, like the nSL statistics (Ferrer-Admetlla *et al.*, 2014) in selscan or the H2/H1 statistics (Garud *et al.*, 2015), although further studies are still needed to understand to what extent hard and soft sweeps can actually be distinguished (Schrider *et al.*, 2015), as well as their relative importance (Messer and Petrov, 2013; Jensen, 2014).

Even hard selective sweeps can be challenging to detect with LD-based statistics (Jensen, 2014). It is advisable to combine several approaches to improve confidence when pinpointing candidate genes for selection. Methods based on LD alone can sometimes miss the actual variants under selection due to the impact of recombination on local polymorphism that can mimic soft or ongoing hard sweeps (Schrider *et al.*, 2015).

All LD-based approaches are more powerful with a relatively high density of markers, such as the ones obtained from whole-genome sequencing, SNP-arrays or high-density RAD-seq, and benefit from using statistics focusing on polymorphism and allele sharing. In a recent study of local adaptation in sticklebacks (Roesti *et al.*, 2015), these statistics have been used on dense RAD-sequencing data to look for recent selection at loci displaying high differentiation (F_ST_). This approach has allowed new candidate loci to be pinpointed, and has confirmed the involvement of those implicated previously (e.g. the *Ectodysplasin* gene). In addition, the identification of large regions displaying high divergence and LD has revealed the importance of large-scale structural variation in shaping genome structure, such as inversions (Roesti *et al.*, 2015).

### Detecting and characterizing selection with the coalescent

If a candidate locus or genomic region has been identified, it is possible to use coalescent simulations to evaluate the strength of selection and estimate the age of alleles. A software such as msms (Ewing and Hermisson, 2010), which is also available in PopGenome, can then be used. However, this requires that population history is known in order to control for other phenomena such as population structure and gene flow. An advantage of full coalescent methods is that they provide a relatively complete picture of the history of individual loci. This can be achieved by modeling coalescence and recombination, and considering variation in mutation rate. However, such methods have long been computationally intensive, and thus difficult to apply to whole genomes. Fortunately, recent computational improvements make their application to whole genomes feasible. A good example is ARGWeaver (Rasmussen *et al.*, 2014), which has allowed candidate genes for long-term balancing selection to be recovered from human data. This method uses ancestral recombination graphs to model the genealogy of each non-recombining block in the genome. Ancestral recombination graphs (ARG) are a generalization of the coalescent and describe the sequence of genealogies along a sample of recombining sequence. Genealogies are estimated for each non-recombining block, and recombination between adjacent blocks is described by breaking the branch leading to the recombining haplotype and allowing it to re-coalesce to the rest of the tree. This succession of local trees joined by recombination events provides a full description of the genealogical history of the data and is therefore a promising approach to characterize positive, purifying or balancing selection while taking into account variation in recombination and mutation rate.

### Identifying variants of functional interest

Characterizing the number of synonymous versus non-synonymous mutations is another approach to detect whether a specific gene is undergoing purifying or positive selection. However, this approach requires an annotated genome. An excess of non-synonymous mutations can signal positive or balancing selection, or a relaxation of selective constraints on a given gene. Annotation of mutations can be done with SNPdat (Doran and Creevey, 2013), or directly in PopGenome, which can also perform tests of selection such as the MK test at the genome scale (McDonald and Kreitman, 1991). Another popular test of selection is the comparison of non-synonymous and synonymous mutations between orthologs from different species, and can be performed in packages such as PAML (Yang, 2007). To recover information about the putative function of a gene or a genomic region, it may be useful to perform a genome ontology (GO) enrichment analysis, using tools such as BLAST2GO (Conesa *et al.*, 2005).

While suggestive, genome scans for selection and association in natural populations cannot be considered as conclusive evidence for the function of a given gene, and need to be combined with functional evidence (Vitti *et al.*, 2013). Such evidence can sometimes be provided by variation in the expression of a candidate gene highlighted by RNA-sequencing data. More often, developmental studies are required, a step that is not always possible for non-model organisms. Pinpointing the exact genetic mutation leading to a change in phenotype is challenging even when combining several tests for selection, and requires whole-genome sequencing data to obtain a near-exhaustive list of mutations. It has been proposed to combine QTL analyses with population genomics to facilitate identification of candidate loci (Stinchcombe and Hoekstra, 2008). Essentially, controlled crosses allow genomic regions associated with a selected phenotype to be identified, while the study of variation in natural populations facilitates the fine-mapping of selected variants in natural populations. However, this requires that the species of interest can be raised in a laboratory or greenhouse, which is unpractical for many research teams. An alternative is the study of candidate genes, for which an extensive description of functional variation is available. For example, in a recent study on passerines (bananaquits), GBS data have been used to obtain a neutral distribution to which patterns of substitution and differentiation were compared at candidate genes for color variation (Uy *et al.*, 2016). Another study on color polymorphism in *Peromyscus* mice used a combination of field experiments, targeted sequencing of candidate genes and neutral regions, and genome-scans for selection. Tests for association between these data were able to show how selection on many mutations at the same locus drive adaptive phenotypic divergence (Linnen *et al.*, 2013).

The combination of tests aimed at different signatures of selection can allow the size of candidate regions to be reduced. For example, combining results from environmental association mapping and genomic scans for selection allows the identification of candidate genes for which a function can be proposed (François *et al.*, 2015). Another common approach relies on the combination of different tests targeting signatures of selection, typically those using the allele frequency spectrum and those using haplotype length. A test of this type has been proposed in human genetics (Grossman *et al.*, 2013), and is called the composite of multiple signals (CMS) test. Nevertheless, signatures of selection can be elusive, and obtaining an exhaustive list of genes under positive selection is unlikely. Further advances will require that methods targeting selection be able to better take into account epistatic interaction and weak selection.

## Suggestions and perspectives

### Estimating selection and demography jointly along a heterogeneous genome

As stated by Lewontin and Krakauer in 1973, “while natural selection will operate differently for each locus and each allele at a locus, the effect of breeding structure is uniform over all loci and all alleles”. Since then, traditional studies on selection have mostly considered that demographic processes act on all loci in the same way across a genome, and that positive selection is mostly rare. This traditional approach has thus tended to disconnect the study of selection from the study of demography (Li *et al.*, 2012).

However, this assumption may be incorrect, and a joint understanding of demography and selection is crucial from this perspective (Figure 3). For example, the large effective population sizes of *Drosophila* have been hypothesized to facilitate a widespread effect of selection across the genome (Sattath *et al.*, 2011; discussion in Li *et al.*, 2012), making both demographic inference and detection of outliers difficult. Other counfounding factors include variation in recombination and mutation rates, and background selection (Ewing and Jensen, 2016), which are difficult to assess with precision in non-model species. Moreover, it has been shown in the last few years that loci involved in reproductive isolation are often also involved in local adaptation. This, combined with variation in introgression rates along the genome, can bias inference about selection and demography (Bierne *et al.*, 2011; Roux *et al.*, 2014). Genomic regions with low recombination rates can lead to reduced polymorphism, and be mistaken for signatures of purifying selection.

These issues can only be addressed by going beyond categorization between methods assigned to either the study of selection or demography, and using the results obtained by one method to inform the other. Such an approach was taken by Tine et al. 2014 in investigating the two different lineages of the European Sea Bass, using a RAD-sequencing approach. Tine et al. took into account variation in recombination rate along the genome to interpret signatures of reduced polymorphism as possibly being the result of selection, low recombination, or a combination of the two (Tine *et al.*, 2014). Since differentiation along the genome seemed to reveal islands resisting gene flow, they could fit a model incorporating variation in introgression rates. This provided improved fit to the data and suggested that islands of differentiation are most likely to be due to locally reduced gene flow after secondary contact. This example illustrates how a combination of descriptive statistics and coalescent analyses can be used to retrieve information from genomic data about both selection and demography.

Most methods do not actually estimate demography and selection jointly, but rather rely on a process where neutral expectations are first drawn from a set of SNPs presumed to be neutral (e.g. intergenic SNPs), followed by a step where the likelihood of a marker being under selection is evaluated. Methods such as BAYPASS or PCAdapt (Table 3) are convenient in both describing population structure and providing preliminary insights into the proportion of loci that do not follow neutral expectations. If this proportion is high, it would suggest recent introgression or an excess of markers displaying high LD (e.g. due to large inversions). However, when this proportion is not too high, outliers can be removed to avoid bias (Schrider *et al.*, 2016) and the remaining loci used to compare neutral models and estimate demographic parameters (e.g. using an ABC framework). These estimated parameters can then be used to simulate sequences or independent SNPs and generate a neutral expectation. Loci that are more likely to be neutral can be used to further calibrate tests for selection such as FLK or BAYPASS (Lotterhos and Whitlock, 2014).

Some recent methods are especially relevant to study both demography and selection at once, while taking into account variation in recombination and mutation rates. For whole-genome data, methods reconstructing ancestral recombination graphs (such as ARGWeaver) have high potential. They allow genealogies to be retrieved along the genome as well as the timing of coalescence events. Such information is ultimately useful for making inferences regarding selection and migration. Recently this method was used in human paleogenomics to quantitatively characterize introgression between modern humans, Neandertals and Denisovans using only a few whole genomes (Kuhlwilm *et al.*, 2016). However, the approach has a high computing and sequencing cost, and is therefore not suitable for studies requiring sampling of many individuals.

Caution must prevail when attempting to apply sophisticated methods to disentangle selection and demography. In a recent review, Cruickshank & Hahn suggest that IMa2, which is commonly used to estimate migration rates, is not able to reliably distinguish between loci resisting gene flow, and those under selection in the absence of gene flow (Cruickshank and Hahn, 2014). In the specific case they highlight (*Oryctolagus cuniculus* rabbits, Sousa *et al.*, 2013), a descriptive statistic that should have captured introgression signatures (d_xy_) did not reveal any evidence for differential gene flow between loci categorized by IMa2. This controversy illustrates that basic description of the data is needed prior using more sophisticated methods. Note however that Cruickshank & Hahn did not address the case of secondary contact, and other methods such as ABC may better detect interruption in gene flow (Sup. Text in Roux *et al*., 2016).

To sum up, the field of population genomics is now moving towards both better integrating the demographic framework in inferences of selection, and, conversely, taking into account selection when reconstructing demographic history. The joint inference of loci under selection and quantification of demographic dynamics is of crucial importance in fields such as landscape genomics or the study of ongoing speciation. It should provide insights into the role of selection, recombination and gene flow in promoting or impairing local adaptation to new habitats. The growing availability of genome-wide data for non-model species is therefore promising, but requires caution and high stringency in our interpretation of observed patterns. With the decreasing cost of sequencing, it has been suggested that NGS will rapidly broaden our perspective on complex evolutionary processes, from biogeography (Lexer *et al.*, 2013) to the genetic basis of traits (Hohenlohe, 2014) and the maintenance of polymorphisms (Hedrick, 2006). While genome heterogeneity in migration, mutation and recombination rates do not necessarily make impossible any conclusion about evolutionary dynamics, they have the potential to blur inferences. The study of DNA sequence variation is already challenging in its own right. Nonetheless, in order to be informative about processes such as selection and demography it should ultimately be combined with other disciplines such as ecology and functional analyses (Habel *et al.*, 2015). This can be done for example by assessing the function of selected genes, the consistency of demographic history with information retrieved from the fossil record or geological history, and the broader integration of population genomics with other fields and methods whenever possible, such as niche modeling, common garden experiments or the study of macro-evolutionary patterns of selection and diversification.

### Beyond SNPs: studying structural variation, transposable elements and epigenetic modifications

Most genome-scale studies of selection and demography have so far focused on SNPs, since they are relatively easy to detect with current technology and their mutation mechanism produces mostly biallelic alleles, making them easier to use for statistical tests. However, many other heritable genetic alterations can affect genomes, including insertions of transposable elements (Villanueva-Cañas *et al.*, 2017), epigenetics modifications such as methylation (Danchin *et al.*, 2011), duplications, inversions, deletions and translocations (Iskow *et al.*, 2012). One of the main issues with this type of variation is that their diversity and their impact on the genome can make them difficult to detect in a systematic way (Iskow *et al.*, 2012), especially for species for which only a draft genome is available. It is however possible to use variation in such genetic alternations to study selection, for example by using differentiation statistics, association to environment or extension of haplotypes. Combining information about variant position and SNP variation in flanking regions is also a powerful way to detect variants under selection (Villanueva-Cañas *et al.*, 2017) as highlighted by a recent study of transposable element insertions in *Drosophila* (Kofler *et al.*, 2012). Recent work also shows that classical summary statistics such as Tajima’s D can be adapted to non-SNP datasets, such as methylations (Wang and Fan, 2014).

Sets of neutral SNPs can be used to control for demography and relatedness between samples when inferring selection. For example, this type of approach has recently been adopted in studies of selection on methylation patterns. In a recent Molecular Ecology issue (Verhoeven *et al.*, 2016), a study using bisulfite precipitation in Valley Oak trees (Gugger *et al.* 2016) was able to place methylated variants associated to climatic variables near to genes known to be involved in response to environment. Another study could show a stronger pattern of Isolation by Distance for methylation-sensitive AFLPs than for regular AFLPs and microsatellites, suggesting a stronger impact of environment on methylation patterns than expected under neutrality (Herrera *et al.*, 2016).

Another potential issue with this type of variation is that there is currently a lack of tools able to simulate their models of mutation, complicating any comparison with neutral models built from SNPs. This is particularly true for transposable elements, for which the assumption of mutation-drift equilibrium is challenging, making comparisons of their allele frequency spectrum with neutral SNPs potentially difficult. For example, a recent burst of transposition can lead to an excess of low frequency elements and recent insertions compared to the expectation under equilibrium, even if transposable elements are not under purifying selection (Bergman and Bensasson, 2007; Blumenstiel *et al.*, 2014). More generally, neutral models would benefit from new ways to model the appearance of genomic variation through time for non-SNP data. This would provide even more conservative assessments of negative and positive selection.

## Acknowledgements

The University of Basel and New York University Abu Dhabi have supported YB’s research in this area. We want to thank two anonymous reviewers, Stephane Boissinot, Joris Bertrand and Anne Roulin for their insightful comments on previous versions of the manuscript.

